# A Microfluidic Chip for LAMP-based Multiplex Detection of Pathogen

**DOI:** 10.1101/2022.05.19.492672

**Authors:** Jingyi Guan, Yunhua Wang, Jing Jin, Guoxia Zheng

**Author notes:** Corresponding Author E-mail addresses.

## Abstract

Early diagnosis of bacterial causing the disease is important for treatment of patent and preventing the spread of pathogen. Utilizing of POCT devices to detect the pathogens on-site will accelerate the diagnosis of infectious disease. By using loop-mediated-amplification, we developed a microfluidic chip for multiplex detection of three bacterial, where the samples were driven by negative pressure were loaded quickly. The performance of the device was preliminarily evaluated. The specificities of the detections were demonstrated. And the LOD for *Escherichia coli, Staphylococcus aureus* and *Pseudomonas aeruginosa* were measured as 17.15, 5.67 and 16.47 ng/μL, respectively. The results demonstrated the feasibility of the method.

---

Infectious diseases are caused by microorganisms, such as bacterial, virus, etc. Traditionally, the techniques, such as microscopy, serology, isolation, selective culturing and biochemistry methods were used to identify the suspect diseases. [1] Culture is considered as gold standard for bacterial diagnosis, while it is labor-intensive and time-consuming. It usually take several days to get results. Early diagnosis of bacterial causing the disease is important for treatment of patent and preventing the spread of pathogen. To allow diagnosis in time, rapid detections are urgently needed. Nucleic acid, including DNA and RNA, are important biomolleculars in the organisms[2]. Each bacterial possess its unique nucleic acid sequence. Amplification of the specific gene of bacterial by PCR (polymerase chain reaction) and more advanced real-time PCR (qPCR) have been developed and widely used for rapid diagnosis of infectious disease [3]. However, these so-called molecular diagnoses depend on trained operator, expensive equipment, which control the temperature reliably and precisely, and operating space. Thus, PCR-based test was mainly performed in central lab. It could not meet the demand of POC testing, which generate faster test results on site and lead to a rapid and reliable diagnosis for infectious disease [4-7].

One alternative strategy to simplify the bacterial detection has been developed. LAMP (loop mediated isothermal amplification) is one of them. As mentioned in name, LAMP amplified the nucleic acids under an isothermal condition. Thus, the expensive thermal cycling equipment is unnecessary. The assay can be done without any sophisticated equipment, without compromising the specificity and sensitivity. Moreover, the LAMP assay was not sensitive to inhibitors original from sample, and tolerate simple pretreatment. They make accurate bacterial identification more rapid and convenient compared to traditional methods [8]. Hence, a great variety of LAMP-based POCT platform were developed.

The second strategy to simplify the device is application of microfluidic. Recently, microfluidic has attracted many attentions. Advantages of microfluidic in developing POCT include less sample and reagent consumption, high-throughput, precise, high sensitivity, reduced detection time and low cost. In nucleic acid detection, the microfluidic could perform many lab processes, including reagent mixing, thermal controlling, biochemical reactions and detection of results. [9-11]

Another option to further simplify the system is reducing the necessity of peripheral pumps, power sources, connectors, and control equipment. For example, capillary pumping in paper based microfluidic analytical system is a class of low cost devices, utilizing capillary force to drive flow of solutions. [12-14] Other examples are degased pumping, finger pumps, and negative pressure pumps, etc. Several negative pressure pumps use a negative pressure chambers as power-source to drive fluid flow. The working channel was separated from negative pressure chamber by a permeability structure. When these devices work, the air in the working channel permeate into negative pressure chamber and then the negative pressure generated in working chamber will actuate the fluid movement [15-17]. However, due to low permeability of currently used structures, the fluid moves slowly. It usually takes several minutes to load the sample.

Here, we reported a new negative pressure actuation structure, where the working channel was connected with negative pressure chamber. Thus, when loading, the sample fills the working channel quickly.

The device consists of two layers of Poly-dimethysiloxane(PDMS). On the top layer, an inlet hole and 12 testing channels (working channel) were designed. The working channels were arranged in a radial manner. In each an inlet channel (Length 15mm × Width 100μm × Height 250μm) connected to a reaction pool (diameter 3mm and height 250μm), where a set of primers corresponding to target bacterial was preloaded. At the end of testing channel, there is a negative pressure chamber(diameter 2mm and height 250μm) to accommodate the residual gas. One test was performed in each testing channel. The sample, mixed with the rest of reagent except for primers, were driven by negative pressure in channel and loaded into the inlet hole and then flow into 12 testing channels.

The device was fabricated by soft lithography as previously described. Briefly, a mask was drawn and printed. SU-8 photoresists (Micro Chem) was spin-coated on silicon wafer and the pattern was transferred. A mixture of Poly-dimethysiloxane (PDMS) monomers (SYLGARD™ 184, Dow Corning, USA) and curing agent, with a ratio of 10:1, was poured on to master wafer. It was then degased and cured. Then, PDMS layer was peeled off. An inlet hole of 1mm diameter was punched at central of the pattern (the junction of testing channel). A PDMS sheet were fabricated and used as bottom layer of the chip. The top layer PDMS was turned upside down. The primer mixture were dripped into reaction pool and air-dried. An irreversible bond between two PDMS layers was created by activating the surfaces with oxygen plasma treatments for 1min. Then the inlet hole was sealed with a PDMS film. A reagent reservoir, which is a PDMS cubic with an invert cone shaped hole in the center, was attached to the film. The device was degased in vacuum tank for 24 h and was sealed in aluminum vacuum packs with a vacuum sealer.

For LAMP (loop-mediated amplification) multiple detection, three bacterial strain, namely, *Escherichia coli* (ATCC25922), *Staphylococcus aureus* (ATCC25923) and *Pseudomonas aeruginosa* (ATCC27853), were used. The bacterial were cultured and collected. The total genomic DNA was extracted by using kit, following the instruction from manufacturer. The targets gene used for these bacterial detections were *ecolc*3109_1[18], *nuc* [19]and *opr* [20], respectively. The primers were designed as previously reported and listed in Table1. Each set of primers include six primers.

**Table 1.**
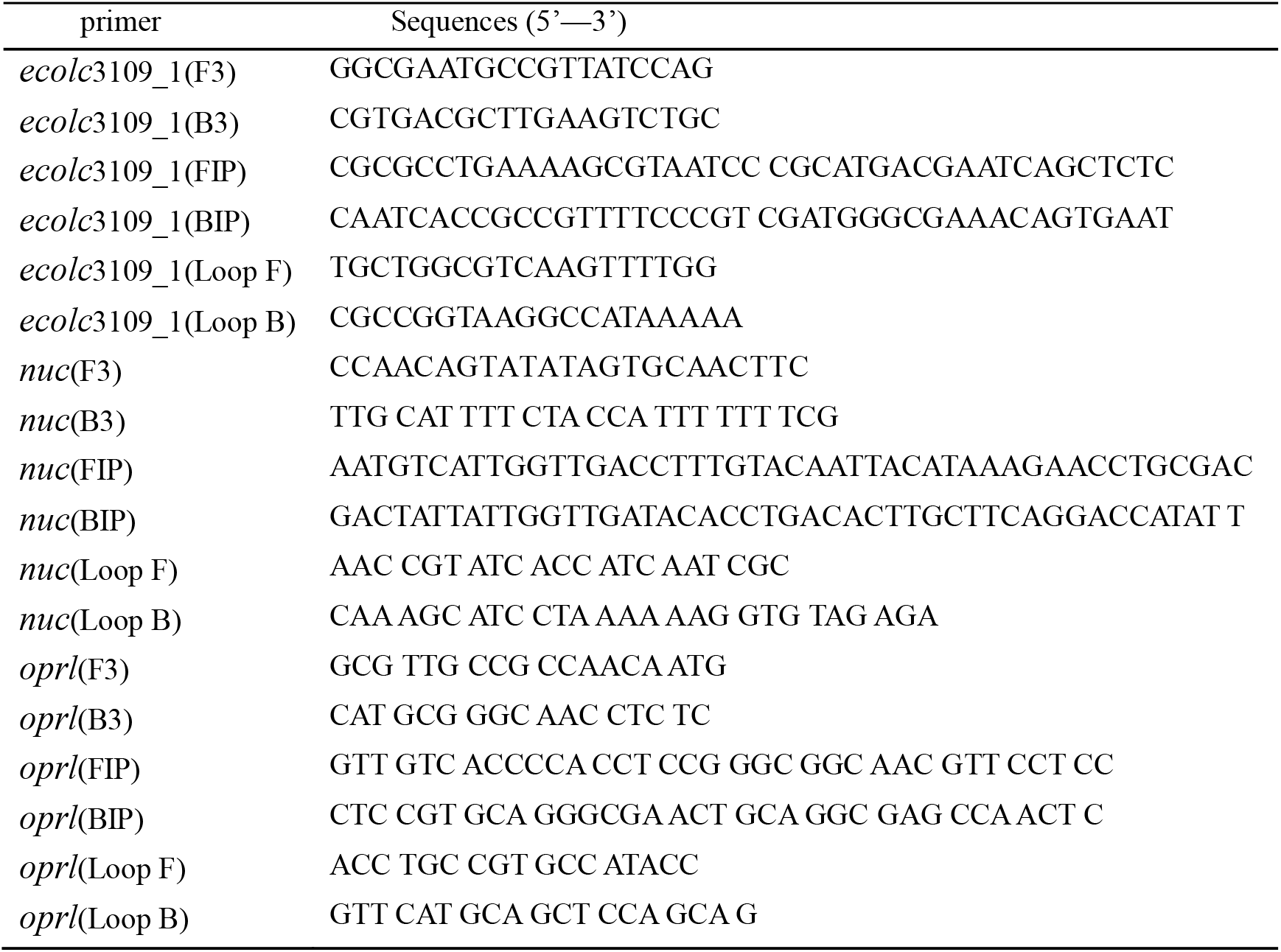
The primer sequences used in this research

The primers of each set were dissolved and mixed with equivalent ratios at final concentration of 10 μmol/L. The mixtures were dropped into corresponding reaction pools, respectively, and air-dried at the room temperature before the chips were assembled.

The LAMP reaction solutions, except for primers, were prepared as follows. Two microliter of 10×ThermoPol reaction buffer (B9004S, NEB, USA), 1.0 μL of Magnesium Sulfate Solution (100mM), 2.5μL of dNTP mixture (10mM each) 1.0 μL of *Bst* DNA polymerase (NEB,USA), 0.5 μL of SYBR Green I and DNA sample were mixed. The ddH2O was added to bring the total volume to 20μL.

The LAMP reaction solution was then dropped into reagent reservoir of degassed device, avoiding generation of air bubbles. A pin was used to cut through the PDMS film at the bottom of reagent reservoir and let the LAMP reaction solution flows in. The solution will be distributed into 12 testing channel equally. When the solution reaches the reaction pool, it will be mixed with pre-loaded primers and initiate amplification if the corresponding template exists. The device was heated in water bath for 1h. The amplified products can be detected by addition of SYBR Green. Then the images of each reaction pool were captured with a fluorescence microscopy equipped with CCD camera.

The testing channels were divided into four groups, for *Escherichia coli, Staphylococcus aureus, Pseudomonas aeruginosa* and negative control assays, respectively. In specificity analyzing, 20 ng DNA of one bacterial was used in each experiment. The results showed that only corresponding bacterial were detected each time and demonstrated high specificity of detection (Figure 2).

**Figure 1.**
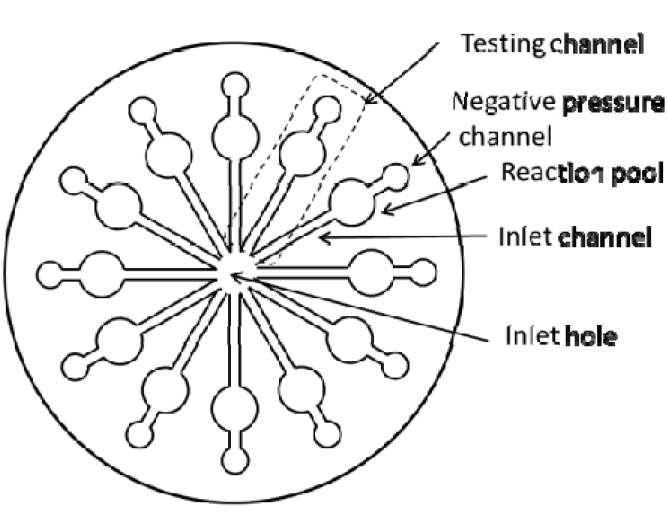
structure of the device.

**Figure 2.**
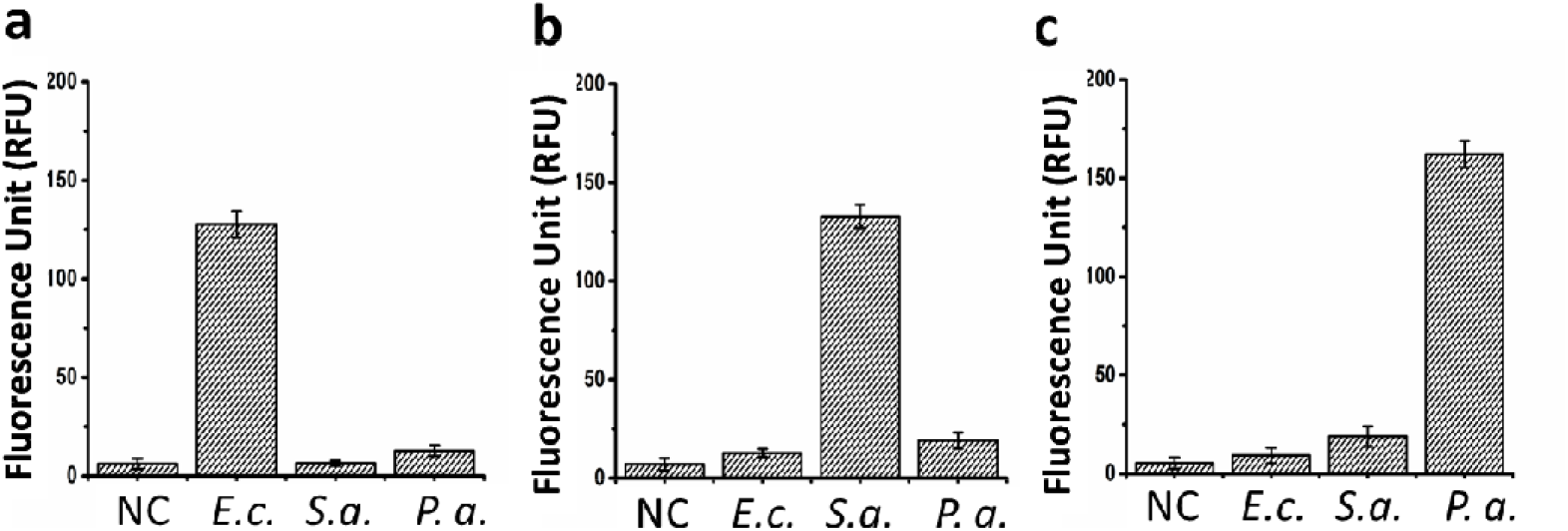
The specificity of the detections. One bacterial genomic DNA was used in each experiment. The results showed that only corresponding bacterial were detected each time and demonstrated high specificity of detection. Note: NC, negative control; *E*.*c. Escherichia coli*; *S*.*a. Staphylococcus aureus* ; *P*.*a*., *Pseudomonas aeruginosa*.

Further we analyzed the interference between reaction pools. The fluorescence images of each reaction pool and its inlet channel were captured. A representative image was presented in Figure 3. The fluorescence signal was detected in the inlet channel, which may due to permeation of amplification production. The signal diffused for about 4mm away from reaction pool. The 15 mm length of inlet channel is long enough to prevent the diffusing of primers to central hole and then to other reaction pools. Thus, the interferences between reactions were avoided by this design.

**Figure 3.**
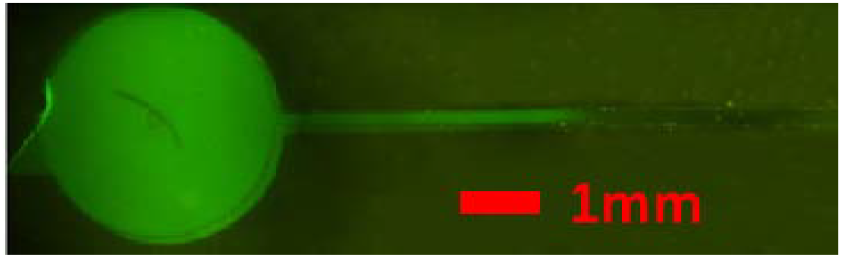
Diffusing of amplification production.

**Figure 4.**
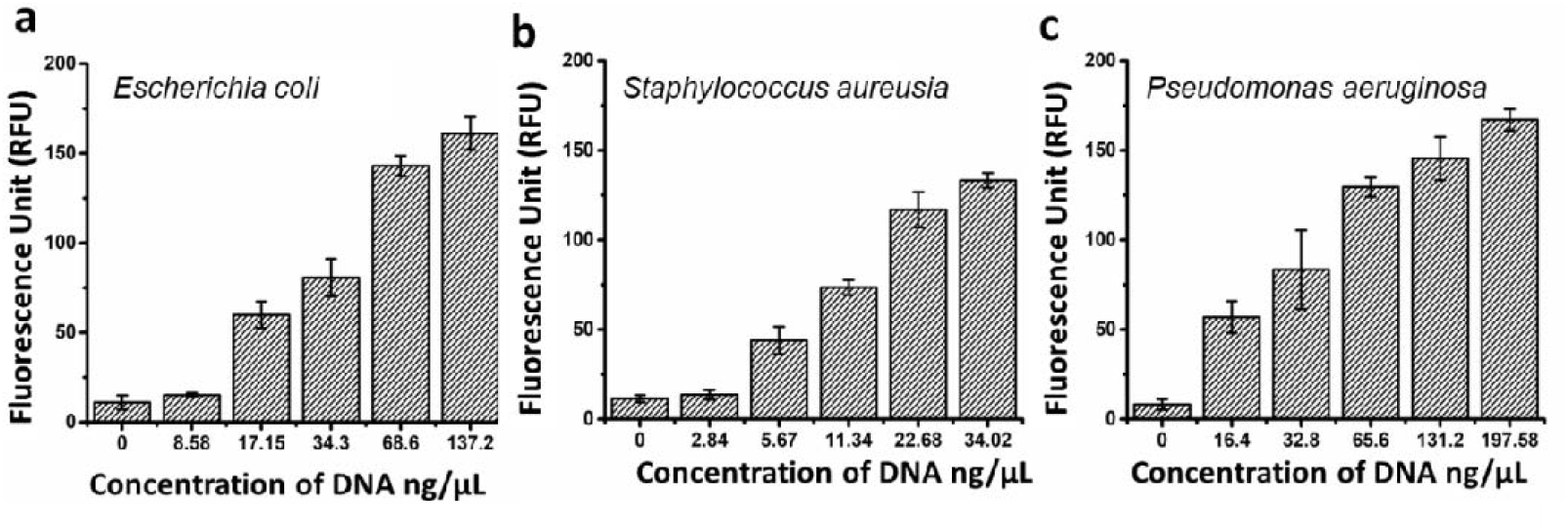
Sensitivity analyzing of three bacterial.

For sensitivity evaluation, serial dilutions of the genomic DNA of each bacterial were used as template. The concentration used in *Escherichia coli* detection were 8.58, 17.15, 34.3, 68.6, 102.90 ng/μL, respectively. They were 2.84, 5.67, 11.34, 22.68 in *Staphylococcus aureus* detection, 34.02 ng/μL, and 16.47, 32.93, 65.86, 131.72 and 197.58 ng/μL in *Pseudomonas aeruginosa*, respectively. Negative controls without primers were included in all assays. In all assays, the fluorescence intensities have linear correlation with the input genomic DNA concentrations. In *Escherichia coli* assay, as shown in Figure 3a, samples with 17.15 ng/μL DNA exhibit a clear fluorescence signal over the threshold value and have a significant difference with the negative control (NC and 8.58 ng/μL DNA. Thus, the LOD to detect *Escherichia coli* was 17.15 ng/μL DNA. Similarly, the LOD of detection to *Staphylococcus aureus* was 5.67ng/μL and to *Pseudomonas aeruginosa* was 16.47ng/μL, respectively. The representative images of detections were shown in top of each figure.

In this study, the LAMP technology was combined with multi-channel microfluidic to develop a device for the simultaneous detection of *Escherichia coli, Staphylococcus aureus* and *Pseudomonas aeruginosa*. It does not require complicated temperature control design, has high sensitivity and specificity, and can detect the DNAs of three kinds of bacteria in time on the spot.

